# Adaptive delayed feedback control disrupts unwanted neuronal oscillations and adjusts to network synchronization dynamics

**DOI:** 10.1101/2022.07.05.498735

**Authors:** Domingos Leite de Castro, Miguel Aroso, A. Pedro Aguiar, David B. Grayden, Paulo Aguiar

## Abstract

Adaptive neuronal stimulation has a strong therapeutic potential for neurological disorders such as Parkinson’s disease and epilepsy. However, standard stimulation protocols mostly rely on continuous open-loop stimulation. We implement here, for the first time in neuronal populations, two different Delayed Feedback Control (DFC) algorithms and assess their efficacy in disrupting unwanted neuronal oscillations. DFC is a well-established closed-loop control technique but its use in neuromodulation has been limited so far to models and computational studies. Leveraging on the high spatiotemporal monitoring capabilities of specialized in vitro platforms, we show that standard DFC in fact worsens the neuronal population oscillatory behaviour and promotes faster bursting, which was never reported in silico. Alternatively, we present adaptive DFC (aDFC) that monitors ongoing oscillation periodicity and self-tunes accordingly. aDFC disrupts collective neuronal oscillations and decreases network synchrony. Furthermore, we show that the intrinsic population dynamics have a strong impact in the susceptibility of networks to neuromodulation. Experimental data was complemented with computer simulations to show how this network controllability might be determined by specific network properties. Overall, these results support aDFC as a better candidate for therapeutic neurostimulation and provide new insights regarding the controllability of neuronal systems.

## 1. Introduction

Brain function relies on coordinated interactions between neurons across a wide range of spatial and temporal scales. However, these interactions might become pathological if excessive synchronization emerges in certain brain areas. This is thought to be at the root of multiple neurological disorders, such as Parkinson’s disease, essential tremor, epilepsy and dystonia^1^. Direct brain stimulation has shown remarkable success in mitigating the symptoms of such diseases when drug therapy is not effective^2^. The interfering electrical stimulation seems to have a desynchronizing effect on the network and disrupt the pathological neuronal oscillations^3,4^. Stimulation is typically applied in an open-loop paradigm, meaning that the stimulus parameters (e.g., pulse frequency, amplitude and duration) are fixed and blind to the ongoing brain activity. This often leads to excessive stimulation and burdensome side-effects^5^. To achieve effective control with minimum side-effects, stimulation should be handled by a closed-loop controller that actuates according to the current state of the brain.

Over the last decades, multiple *in silico* studies focused on developing closed-loop stimulation protocols to desynchronize pathological oscillations in computational models of neurological disorders^6–11^. However, these have rarely translated into studies with real biological neuronal systems. A recent exception is the implementation of a closed-loop controller, previously presented in an *in silico* study^10^, that used calcium imaging feedback to desynchronize neurons in brain slices^12^. However, instead of disrupting the underlying oscillation, this method spread the oscillation through different out-of-phase neuronal clusters.

A particularly prevalent method in the field is Delayed Feedback Control (DFC), a technique from chaos theory developed to stabilize/destabilize chaotic systems^13,14^. DFC controls the system by applying a feedback signal proportional to the difference between the current oscillation amplitude and the oscillation amplitude from a fixed delay into the past. The choice of delay determines whether the controller enhances or suppresses the oscillations. This method has interesting properties that make it attractive for neuromodulation applications: it does not require prior knowledge about the controlled system, the actuation signal vanishes as the system approaches the desired trajectory (be it a stable orbit or its suppression), it is simple and easy to implement both in software and hardware and the eventual technical latencies can be compensated by adjusting the delay accordingly. DFC has been extensively applied in the context of neuroscience, with studies focusing on specific disorders such as Parkinson’s Disease^15–20^ and epilepsy^21^. However, there is still controversy in the literature regarding its efficacy and whether it may instead promote synchrony depending on the baseline levels of synchrony of the modelled network^22^.

To the best of our knowledge, DCF has only been used to control neuronal systems *in silico*. Computational models are undoubtedly useful to test different hypothesis in a controlled environment; nonetheless, they are still an oversimplified representation of real neuronal systems. Biological neuronal networks display complex time-changing dynamics that cannot yet be fully captured with computational models. Also, the effects of electrical stimulation on the overall network dynamics are far from being fully understood, much less simulated. Therefore, testing DFC with biological neuronal networks that display synchronized oscillatory firing is a fundamental step to assess its viability as a brain stimulation protocol for the treatment of neurological disorders.

Neuronal cultures on microelectrode arrays (MEAs) are useful a testbed for this task: after several days *in vitro*, these cultures self-organize into complex interconnected networks with quasiperiodic synchronized bursting. MEAs enable full control of the neuronal activity, providing concurrent recording and stimulation at multiple points of the population with high spatial and temporal resolution. This versatility fostered several studies aiming to control specific features of neuronal activity with closed-loop electrical stimulation in MEAs, such as the mean population firing rate using a proportional controller^23^, the network response probability and latency to stimulation using a proportional-integral controller^24^ and response strength to stimulation using reinforcement learning^25,26^.

In this work, we implemented, for the first time, the DFC algorithm to disrupt the synchronized bursting of biological neuronal populations. The controller modulates the frequency at which individual electrical pulses are sent. We opted for such a pulsatile actuation signal instead of a continuous signal to mimic the currently available stimulators used in implantable devices. The cells were cultured on MEAs and the network activity was controlled in real time with very low latencies using specialized hardware and custom open-source software. We show that traditional DFC lacks the adaptability to cope with the malleable dynamics of biological networks and may actually impose a new firing periodicity. As an alternative, we developed and implemented an adaptive DFC (aDFC) algorithm that self-tunes to better predict the current oscillation phase. We show that aDFC reduces the synchronization of the network and efficiently disrupts the baseline oscillation. We identified and characterized a new feature of neuronal networks that should be taken in consideration in future studies: self-organized neuronal cultures have different propensities for modulation conditioned by their intrinsic population dynamics. Using computer simulations, we show that this controllability can be conditioned by specific parameters of the network architecture, namely the excitatory/inhibitory balance and overall synaptic weights of the network connections. Finally, we show that in a network exhibiting spontaneous transitions to asynchronous firing, aDFC (but not DFC or random stimulation) can induce and sustain this firing regime.

## 2. Results

To assess the potential of DFC as a closed-loop brain stimulation protocol (Figure 1.A, top) for suppressing pathological neuronal oscillations, we developed an *in vitro* setup composed of hippocampal networks cultured on microelectrode arrays (MEAs), a workstation running the control algorithm and specialized hardware (2100MEA-System, Multichannel Systems) interfacing both environments at very low latencies (Figure 1.A, bottom). Our *in vitro* setup provides a powerful testbed to analyse DFC algorithms as it incorporates neuronal populations that display synchronous oscillations (here in the form of quasiperiodic network bursts with periodicities in the seconds range), an acquisition system that measures the population’s extracellular activity at high spatiotemporal resolution enabling the inference of synchrony among neurons, on-demand electrical pulse generators to stimulate the network at high spatiotemporal resolution and a feedback loop with low actuation latency in the milliseconds range (orders of magnitude below that of the neuronal dynamics under control).

**Figure 1.**
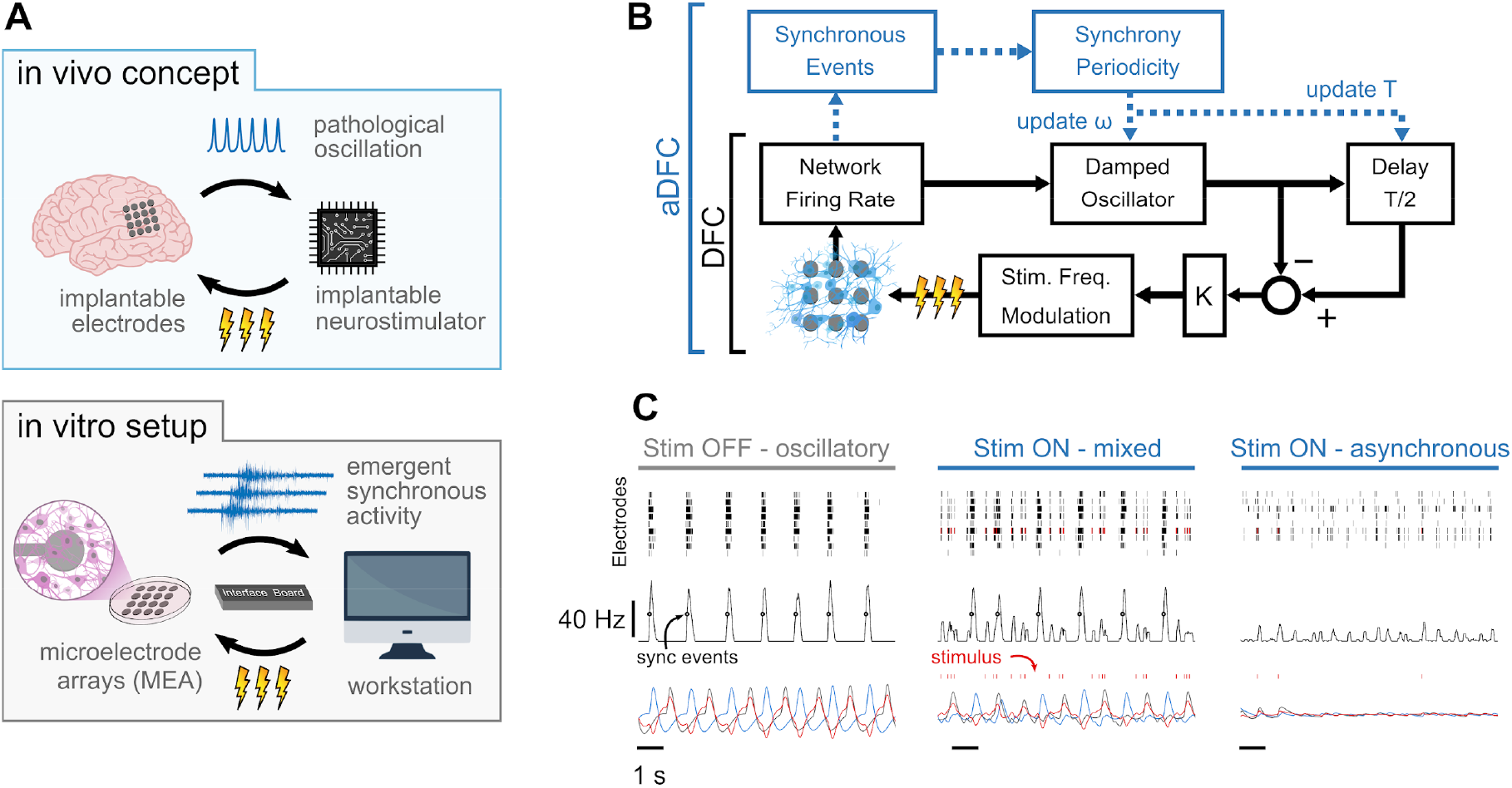
Controlling pathological oscillations with Adaptive Delayed Feedback Control (aDFC). (A) *In vivo* concept of implantable closed-loop neurostimulation and the analogous *in vitro* setup developed. Here, we have dissociated hippocampal neurons as a brain, microelectrode arrays as implantable electrodes, emergent periodic network bursting as pathological synchronous oscillations, a workstation as an implantable chip and an external stimulator as an implantable neurostimulator. (B) Scheme of the Delayed Feedback Control algorithms. The instantaneous population firing rate is calculated and filtered by a damped oscillator. The actuation signal (stimulation frequency) is obtained by subtracting the ongoing oscillation to the past half-cycle delayed oscillation. The natural frequency of the oscillator, ω, and the duration of the half-cycle delay are updated online (adaptive component in blue) by detecting the synchronous events and their current periodicity, T. (C) Controller computations under different firing regimes. Spike detection is performed for the monitoring electrodes (top) and the spike train is convolved online with a square window to compute the instantaneous population firing rate (middle). Synchronous events (black circles) are detected using a threshold crossing method and used to update the oscillation periodicity. The actuation signal (red) is built by subtracting the oscillator and delayed oscillator signals (blue and grey, respectively) and translated to a stimulation frequency signal.

The core of the DFC algorithm used (Figure 1.B, black) is analogous to implementations reported in the literature (see Methods section for details). After spike detection (Figure 1.C, top row), the instantaneous population firing rate is calculated (Figure 1.C, middle row) and filtered by a damped oscillator (Figure 1.C, bottom row in blue). The resulting signal is a good representation of the oscillatory behaviour of the network, such as that exhibited while the stimulation is turned OFF (Figure 1.C, left). The feedback signal (Figure 1.C, left, bottom in red) is then created by subtracting the ongoing oscillation to the half-cycle delayed oscillation (Figure 1.C, bottom row in grey). This way, when the activity is oscillatory, the actuation signal is maximal at the antiphase of the neuronal oscillation. The obtained feedback signal is linearly translated to stimulation frequency by applying a fixed gain. Since the actuation signal is explicitly dependent upon the neuronal oscillation, if this firing pattern vanishes, the actuation signal decays to 0 and electrical stimuli are no longer sent (Figure 1.C, right). Expanding on the DFC algorithm, and to respond to the dynamic nature of biological neuronal systems, we improved canonical DFC to adaptive DFC (aDFC) by adding an extra component that detects the synchronous firing events (Figure 1.C, black circles in middle row). This way, aDFC monitors the current periodicity of the neuronal oscillation (T) and updates the delay duration (T/2) and natural frequency of the oscillator (ω) accordingly (Figure 1.B, blue and Figure S1).

The neuromodulation performance of the control algorithms was analysed in a series of experiments with multiple neuronal populations on MEAs. Each experiment had three stages: 5 minutes of spontaneous activity pre-stimulation (OFF), 5 minutes under stimulation (ON) and 5 minutes of spontaneous activity post-stimulation (OFF). Every network, here defined as a neuronal culture on a given day *in vitro*, was submitted to three different stimulation algorithms: standard DFC, aDFC and random stimulation. During random stimulation, the electrical pulses were triggered by a Poisson process with an average frequency identical to that obtained in the preceding aDFC test. In all algorithms, stimulation was provided as monophasic negative voltage pulses at controlled timings. To ensure that the individual pulses had a comparable effect on the different neuronal networks, we chose a single stimulation electrode that triggered a neuronal response measured by neighbouring electrodes and the minimum voltage amplitude that achieved that response (see Methods section).

### 2.1 Adaptive component is required due to the dynamic nature of neuronal populations

Changes in the neuronal dynamics caused by the stimulation protocols can be evaluated using the wavelet transform of the instantaneous network firing rate (Figure 2.A and 2.B). This representation evidences how the bursting periodicity of the network changes over time. Consistent inter-burst intervals (i.e., consistent neuronal oscillations) will appear as a low frequency horizontal line. When stimulation was activated, both DFC and aDFC showed clear effects on the neuronal dynamics. DFC stimulation imposed a change to a faster and very consistent oscillation (DFC periods in Figure 2.A and C). This happens because DFC initially forces the network to fire in its antiphase, creating a change in the bursting periodicity. Since this change is not monitored to update the controller’s delay, the delayed oscillator (Figure 1.C grey) no longer matches the anti-phase of the ongoing oscillation. Therefore, stimulation becomes prejudicial as it is sent in phase with the new oscillation, reinforcing it (DFC period in Figure 2.C).

**Figure 2.**
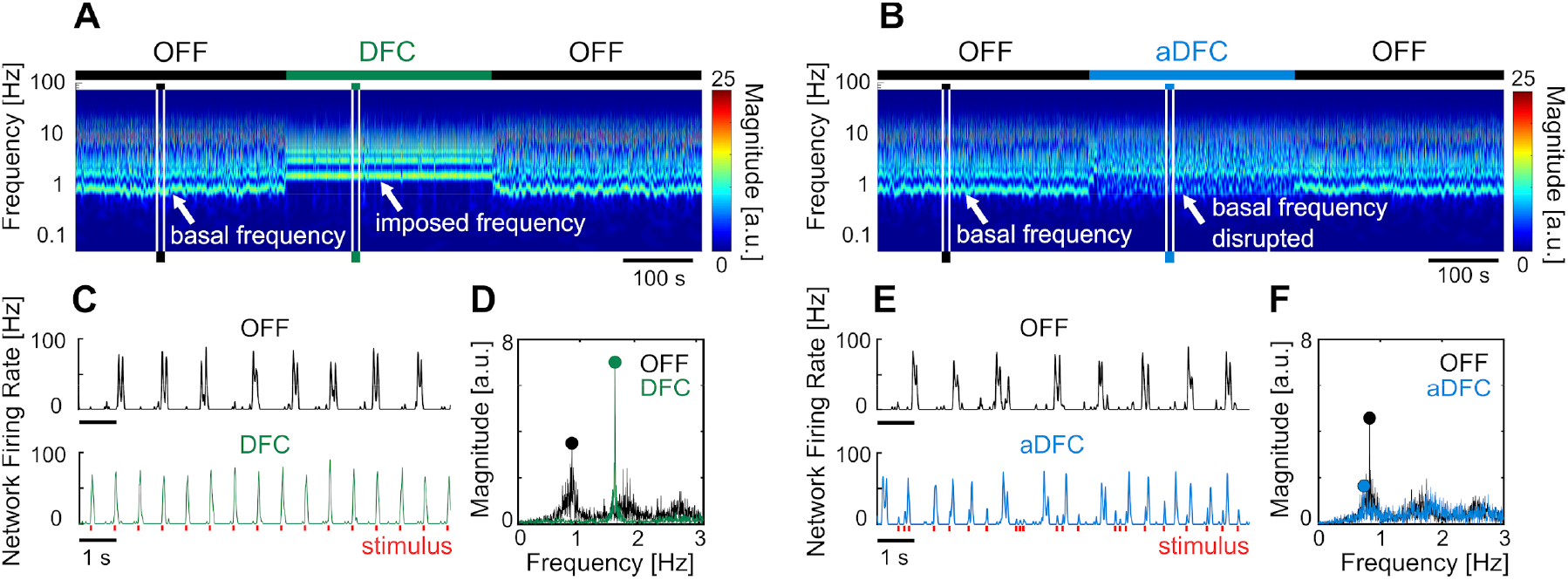
Representative example of the effect of DFC and aDFC stimulation protocols. (A,B) Wavelet transform of the instantaneous firing rate of a network under DFC (A) and aDFC (B) protocols. (C,E) Example of the instantaneous firing rate of the network before stimulation (black) and during DFC (C, green) and aDFC (E, blue) stimulation. The signals showed in C and E correspond to the narrow time windows marked in A and B, respectively. (D,F) Discrete Fourier transform of the instantaneous firing rate of the network before stimulation (black) and during DFC (D, green) and aDFC (F, blue) stimulation. The main oscillation under each condition is marked with a coloured circle.

On the other hand, aDFC disrupts the spontaneous oscillation frequency exhibited by the network, creating a firing regime where there is no predominant oscillation (aDFC periods in Figure 2.B and 2.E). In this case, the controller is constantly forcing the network to fire in antiphase and aDFC adapts as the network oscillation changes with the stimulation. To quantify this effect, we calculated the magnitude and frequency of the main oscillation by extracting the maximum of the discrete Fourier transform of the instantaneous firing rate for the ON and OFF segments (Figure 2.D and F).

### 2.2 Network controllability is conditioned by intrinsic population dynamics

The effect of the different stimulation protocols was evaluated using four metrics calculated for the OFF and ON periods: synchrony, firing rate, oscillation frequency and oscillation intensity. Synchrony measures the degree of co-activation of multiple electrodes (see Methods); firing rate corresponds to the average number of spikes per electrode per unit of time; oscillation frequency and intensity are, respectively, the frequency and amplitude of the main peak of the Fourier transform (Figures 2.D and F).

The three stimulation protocols were repeated multiple times for each network following the OFF-ON-OFF routine. Their effects on a given network are evidenced by normalizing the values of each feature to the initial OFF period (before stimulation) across the multiple trials (Figure 3.A). This also reveals how the dynamics are recovered once the stimulation is turned OFF.

**Figure 3.**
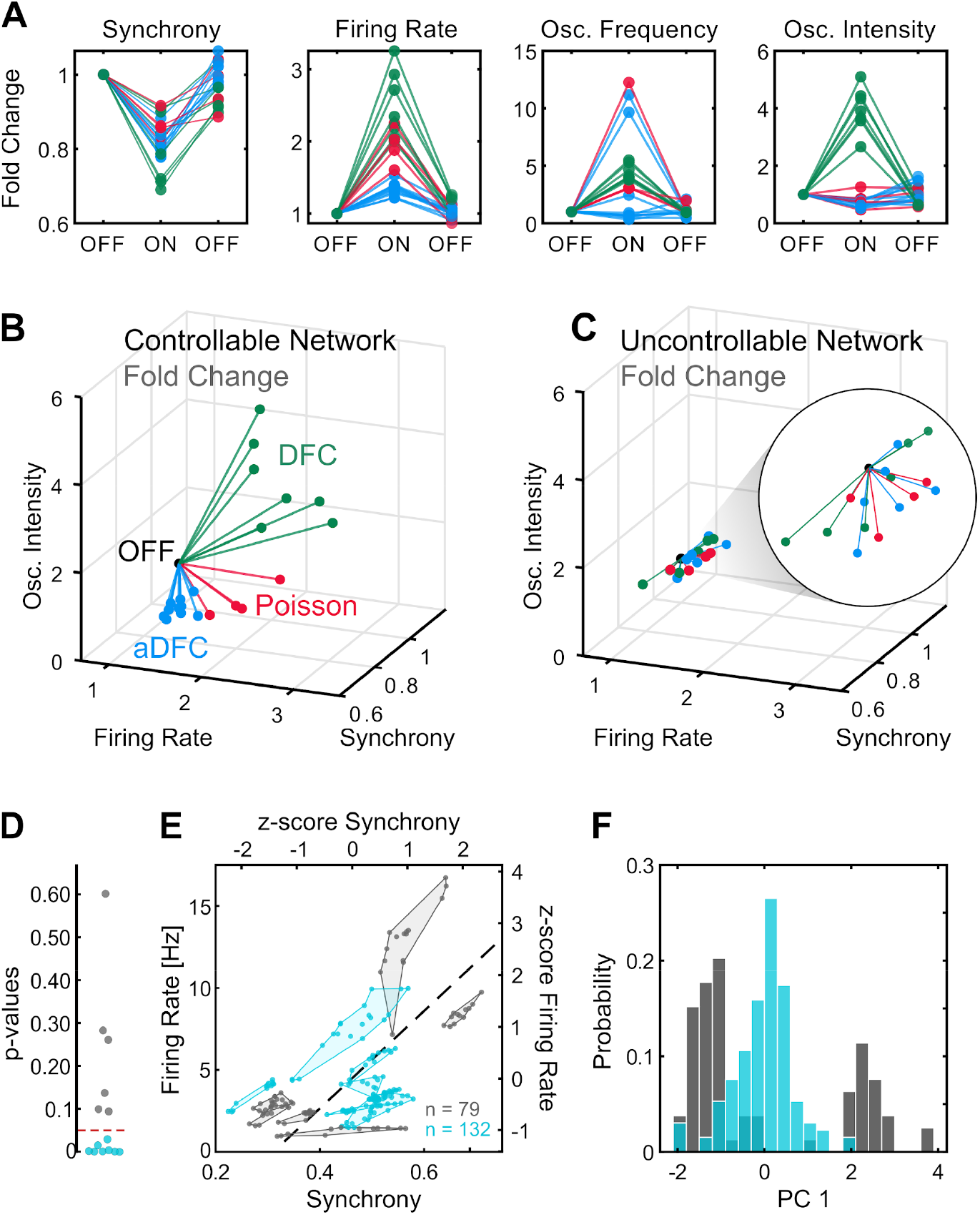
Comparison of controllable and uncontrollable networks. (A) Results obtained from multiple repetitions of the three stimulation protocols for a single, representative network. We calculated the synchrony, firing rate, oscillation frequency and oscillation intensity for the three segments – pre-stimulation (OFF), stimulation (ON), post-stimulation (OFF) – of each experiment and normalized each metric to the value of the pre-stimulation period. Each triplet of points corresponds to one experiment performed with either aDFC (blue), DFC (green) or Poisson (red) stimulation. (B) Three-dimensional (3D) feature space – Firing Rate vs. Synchrony vs. Oscillation Intensity – for the network in A. Each line corresponds to the metrics fold change from the initial OFF to ON (from the first to the second point in A). This network is considered controllable since each protocol consistently drove the network to a distinct subspace, meaning that it had a unique and reproducible effect. (C) Representative example of the 3D feature space for an uncontrollable network where the effect of each stimulation protocol is indistinguishable from the others. (D) Networks were separated into controllable (blue) and uncontrollable (grey) by computing the MANOVA of the ON metrics for the different stimulation protocols. Networks with p-values lower than 0.05 had separable effects in the 4D feature space of the metrics and were thus considered controllable (8/14). (E) Pre-stimulation firing rate and synchrony for all experiments performed with each network divided into controllable and uncontrollable networks. Each dot corresponds to one experiment performed with either DFC, aDFC or Poisson in a given network. The experiments performed with the same network are confined in a polygon. The black traced line corresponds to the first principal component (PC1) of the standardized synchrony and firing rate defining a relevant descriptor of the neuronal dynamics. (F) Distribution of the baseline neuronal dynamics along PC1, defined in E, showing that the controllable networks have an intermediate baseline level of firing rate and synchrony.

Some networks were consistently driven to distinct regions of the features space with each stimulation protocol (Figure 3.B). We refer to these as *controllable* networks – hypothetically, an ideal controller could lead such networks to any desired reachable state. On the other hand, some networks did not exhibit consistent modulation when stimulated using the different protocols (Figure 3.C). We termed these *uncontrollable* networks. Based on this definition, each network was classified as either controllable or uncontrollable by calculating the multivariate ANOVA for the ON values of the four metrics (see Methods for details). Networks with low p-values (p < 0.05) (i.e., networks with significantly distinct neuromodulation results for the different stimulation protocols) where considered controllable (Figure 3.D).

Having recognized that some networks are controllable, we questioned if there are any common traits that make them more prone to neuromodulation. Even though there may be an indefinite number of hidden variables that influence the controllability of a neuronal population, some of these properties can be translated into specific firing dynamics, which we can access with the MEAs. If that is the case, then we expect the controllable and uncontrollable networks to have distinct fingerprints in their spontaneous firing dynamics. To evaluate this hypothesis, as well as ensure that the uncontrollable networks did not simply result from an ineffective stimulation electrode, we compared the spontaneous firing rate and synchrony during the initial OFF period (before stimulation) for all the experiments performed with all networks. This revealed that, as hypothesised, controllable and uncontrollable networks lie in different regions of the feature space (Figure 3.E and Figure S2). Furthermore, controllable networks are in the centre, which is associated with intermediate levels of spontaneous firing rate and synchrony. This can be further evidenced by calculating the principal components (using Principal Components Analysis, PCA) of the standardized map and projecting the values of each experiment along the principal component. A histogram shows that the experiments performed with controllable networks are in the centre, surrounded by a bimodal distribution of the experiments performed with uncontrollable networks (Figure 3.F).

### 2.3 aDFC disrupts oscillations and decreases overall synchrony in controllable networks

The performances of DFC, aDFC and Poisson stimulation were evaluated by comparing their average effects on each controllable network (Figure 4). Uncontrollable networks are not considered here as their results are, by definition, not reproducible. We included trials without stimulation for each network as a control group to capture the natural variability of each metric; in these, we calculated the metrics for two consecutive 5 min blocks of spontaneous activity, representing the initial OFF and ON periods.

**Figure 4.**
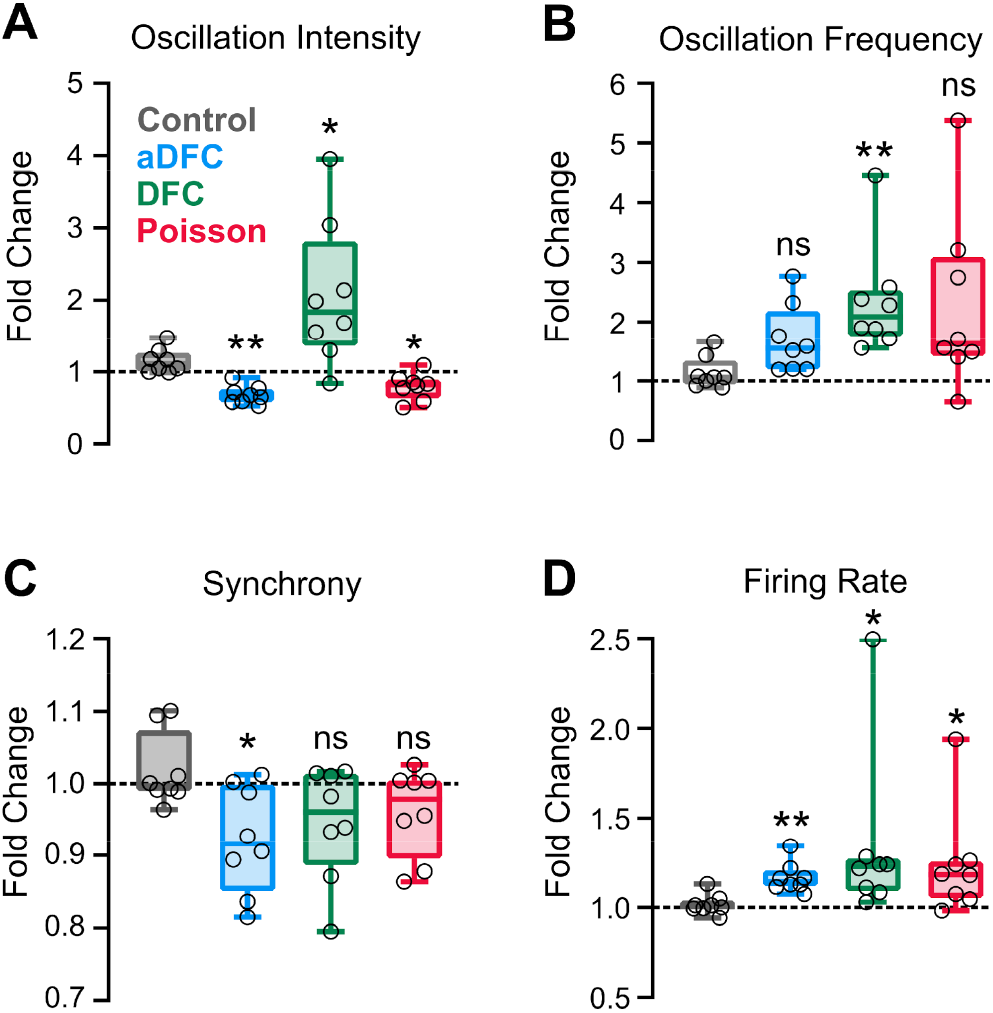
Effect of aDFC, DFC and Poisson stimulation in controllable networks. Fold change in oscillation intensity (A), oscillation frequency (B), synchrony (C) and firing rate (D) of the 5 min stimulation period compared to the 5 min pre-stimulation period with aDFC (blue), DFC (green) and Poisson (red) stimulation. The control group (grey) corresponds to baseline recordings where there is no stimulation and thus portrays the natural variability of each metric within two subsequent 5 min blocks. Each point represents the average effect of a stimulation protocol across all trials for a given network. We used t-tests to compare each group with the control (n = 8; * p < 0.05; ** p < 0.01; details of statistical tests in Table S1)

The most significant distinction between control algorithms was evidenced at the level of the bursting oscillations. As already demonstrated for a representative example (Figure 2), standard DFC led to more consistent oscillations (Figure 4.A), typically doubling the baseline frequency (Figure 4.B, green). This erratic behaviour did not occur with aDFC, as the intensity of the bursting oscillation was significantly decreased (Figure 4.A, blue). Random stimulation with the same stimulation frequency led to a wide range of bursting oscillation frequencies (Figure 4.B, red) but with low oscillation consistency (Figure 4.A, red). The only stimulation protocol that led to a significant decrease in synchrony was aDFC (Figure 4.C). All methods led to an increase of the average firing rate (Figure 4.D), which can be attributed to the fact that excitatory input was always provided and with similar average stimulation frequency for the three methods (Figure S3).

### 2.4 Controllability is affected by synaptic strength and balance between excitation and inhibition

We then sought to understand whether the controllability subspace found for the neuronal cultures portrayed any fundamental property of these type of systems and what parameters could govern such behaviour. We modelled randomly connected networks of 1000 Izhikevich neurons^27^ with different proportions of regular spiking excitatory neurons and fast spiking inhibitory neurons. We also varied the overall synaptic weight (see Methods for details). Each condition was simulated five times. The simulations were composed of two different periods: spontaneous activity (OFF) and controlled activity (aDFC ON). For each simulation, we computed the overall synchrony and firing rate for the OFF (Figure 5.A) and aDFC ON periods (Figure 5.B). Low percentages of excitatory neurons and weak synaptic weights within the network led to sparse and chaotic activity (regions 1 in Figures 5.A-C). In this scenario, the controller is sporadically activated due to fluctuations of neuronal activity, creating brief moments of synchronization (Figure 5.C, top). High percentages of excitatory neurons and strong synaptic weights led to highly ordered network dynamics with strong bursting activity (regions 3 in Figures 5.A-C). Here, the controller could not disrupt the dominating spontaneous activity (Figure 5.C, bottom). However, in the transition between these two network states, there was a parameter subspace that generated intermediate baseline levels of firing rate and synchrony (regions 2 in Figure 5.A-C). In parallel to what was found *in vitro*, the networks were controllable in this intermediate region. Here, aDFC could desynchronize the network shortly after activating the stimulation (Figure 5.C, middle). Thus, among the multiple possible parameters governing network controllability, the excitatory/inhibitory balance and overall network connectivity strength could explain the separation into controllable and uncontrollable networks seen *in vitro*.

**Figure 5.**
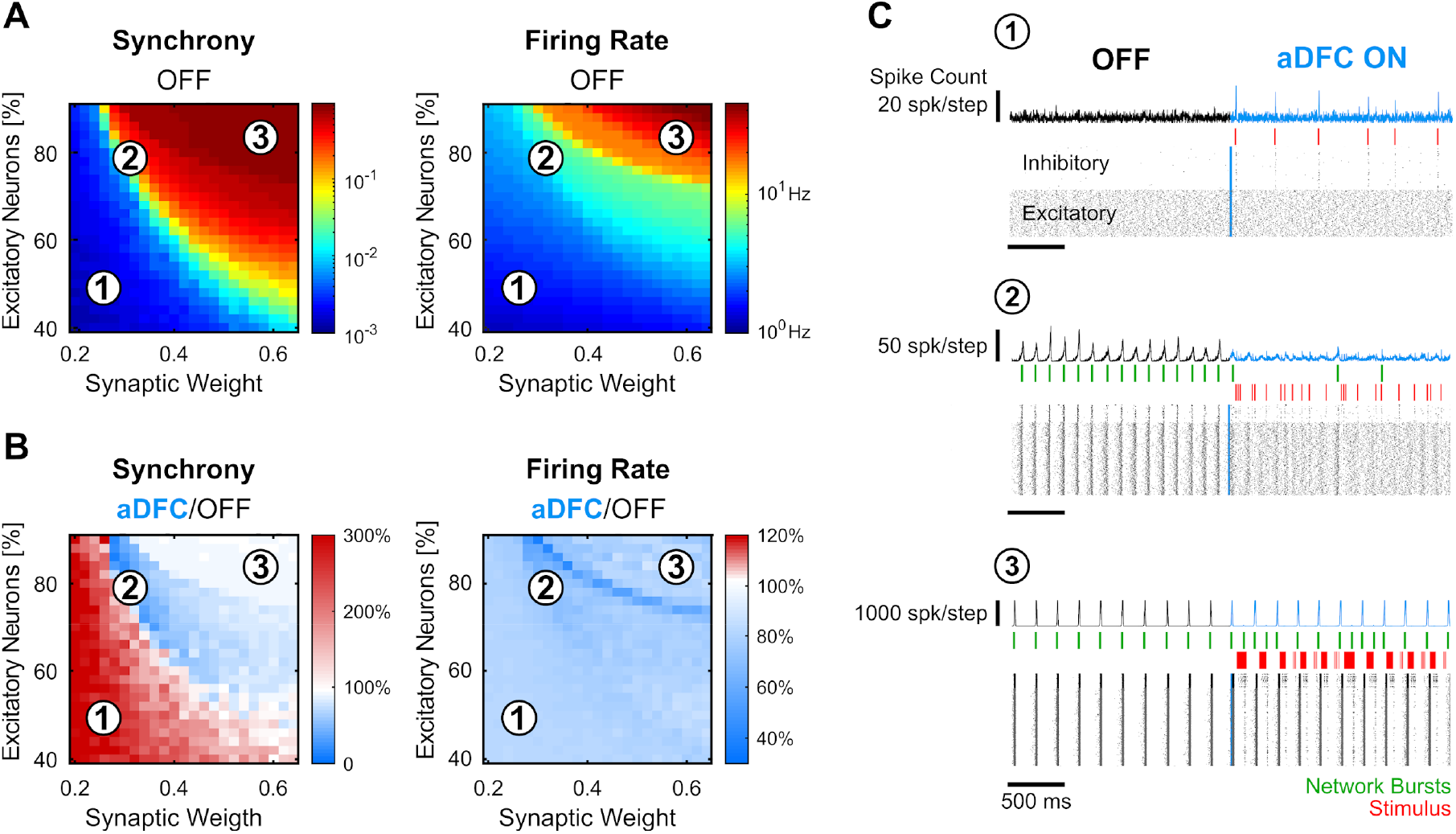
*In silico* networks also present a controllable subspace for intermediate baseline levels of firing rate and synchrony. (A) Level of synchrony (left) and firing rate (right) for the period of spontaneous activity. Each pixel represents the average synchrony and firing rate of five different simulations under the same conditions of excitatory/inhibitory balance and synaptic weight. (B) Average change in synchrony (left) and firing rate (right) of aDFC ON period compared to the OFF period. (C) Representative example of the simulations under three distinct network parameterizations (1, 2 and 3 in A and B). Each panel has the instant spike count at the top, the raster plot at the bottom and the network bursts (green) and triggered stimuli (red) in the middle.

### 2.5 aDFC can promote transitions to stable asynchronous states

The majority of the *in vitro* cultures had monostable dynamics, presenting a very consistent firing pattern over time (e.g., OFF periods of Figure 2). In these networks, even though aDFC reduced neuronal synchronization (Figure 4.C) and disrupted baseline oscillatory activity (Figure 2.D-F and Figure 4.A), it did so by creating sparser bursting with no particular periodicity. In monostable networks, aDFC did not promote a phase transition to a stable asynchronous state as opposed to what is typically seen in DFC computational studies.

We also found a less common type of networks that had multi-stable dynamics, spontaneously transitioning between different firing regimes. Multi-stable networks were particularly interesting when they presented sporadic transitions to an asynchronous state (AS) as it showed that these networks could autonomously sustain asynchronous activity, even if only for short periods of time (Figure 6). The different stable states were identified using unsupervised clustering on the values of firing rate and synchrony calculated over a sliding window (Figure 6.A,B). A given state was considered asynchronous if its average synchrony was below 0.5 and the mean firing rate was below the average of that network. The obtained clusters correlate with the different regimes seen in the frequency domain (Figure 6.C).

**Figure 6.**
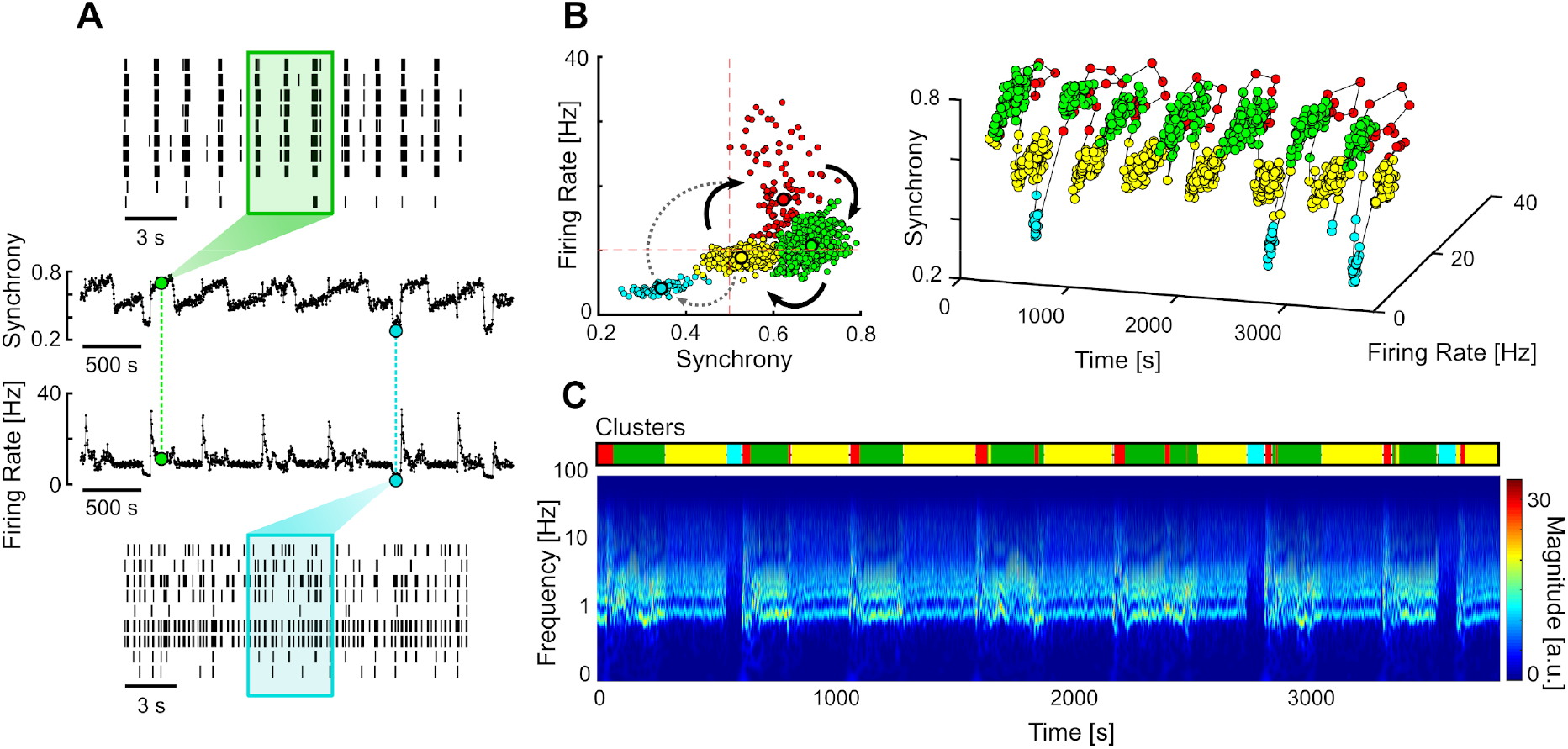
Multi-stable network with sporadic transitions to asynchronous state. (A) Spontaneous synchrony and firing rate over time (middle). Representative raster plots of the synchronous (top) and asynchronous states (bottom). The coloured boxes represent the sliding window over which the firing rate and synchrony are calculated. (B) Synchrony vs. Firing Rate phase plot evidencing the cyclic behaviour of the network, with sporadic transitions to the asynchronous state (blue cluster). The clusters were identified using unsupervised Gaussian mixture models. (C) Changes in the frequency domain correlate with the automatically identified clusters (top bar)

To evaluate the effectiveness of each stimulation protocol in promoting transitions to AS, we compared the percentage of time spent in AS during the OFF and ON periods in multi-stable networks (Figure 6). Since our experimental protocol was composed of 5 min blocks (OFF-ON-OFF), the desynchronizing effect of the closed-loop stimulation was only clear for networks that spontaneously exhibited rare and brief (considerably less than 5 mins) AS. This way, a significant increase in the time spent in AS during the ON periods was most likely caused by the closed-loop stimulation. Out of all the networks tested, one presented the required criteria to evaluate this effect (Figure 7.A and B).

**Figure 7.**
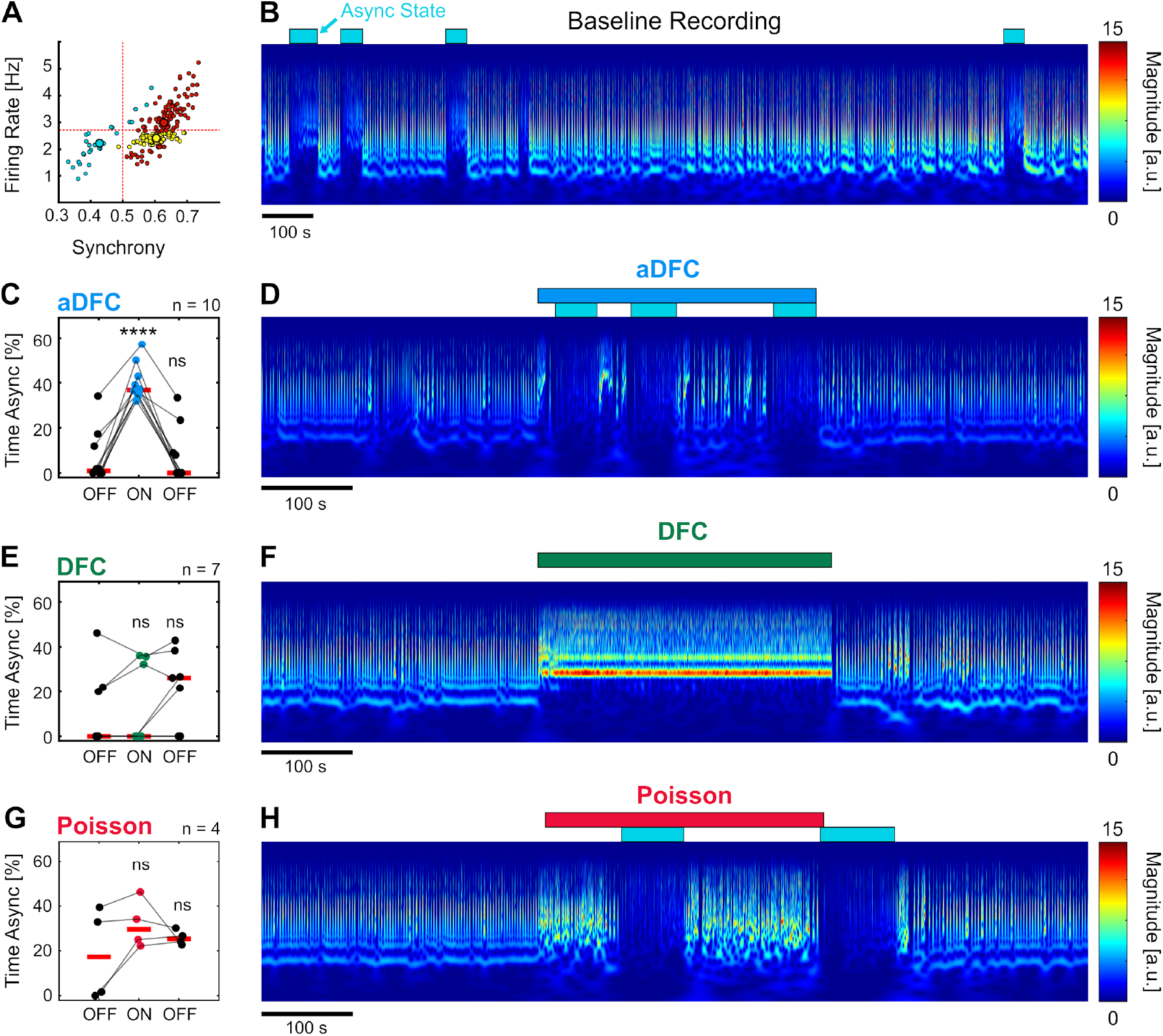
aDFC can promote transition to asynchronous state. (A) States of spontaneous activity for a given multi-stable network. Blue cluster represents the (sporadically reached) asynchronous state. (B) Frequency domain of the recording considered in A. Blue rectangles represent the regions associated with the automatically identified asynchronous state. (C, E, G) Percentage of time in asynchronous state during the OFF-ON-OFF protocols for aDFC (C), DFC (E) and Poisson (G) stimulation. Each triplet of points represents an experiment performed with the corresponding stimulation protocol. We compared the time spent in AS during the ON and the second OFF periods with the initial OFF period using t-tests (**** p< 0.0001; details of statistical tests in Table S2). (D, F, H) Representative example of an experiment performed with aDFC (D), DFC (F) and Poisson (H) stimulation portrayed in the frequency domain. The stimulation period is represented with the associated coloured bar on the top. The identified asynchronous states are represented by the blue rectangles, as in B.

aDFC consistently promoted desynchronization in all the trials performed with this network, leading to a considerable increase of time spent in AS (Figure 7.C,D). DFC stimulation, on the other hand, tended to impose a new bursting periodicity (as already reported), sometimes interrupted by transitions to AS (Figure 7.E,F). However, the percentage of time spent in AS during DFC stimulation is within the range of spontaneous transitions and thus cannot be directly attributed to DFC. Random Poisson stimulation with identical stimulation frequency promoted sparser bursting with some transitions to AS, but these are also within the range of spontaneous transitions (Figure 7.G,H). This means that the precisely timed stimuli determined by aDFC are causing the transitions to the asynchronous state.

## 3. Discussion

Controlling brain activity is a long standing goal in neuroengineering and holds great potential for multiple clinical applications. This is challenging because neuronal systems are inherently complex, with non-linear (arguably chaotic) dynamics, virtually impossible to fully observe, have limited access for an external controller to actuate and the nature of the actuation itself is highly restricted. Most of these constraints are circumvented in computational studies, allowing researchers to explore solutions in a highly controlled environment. However, under such simplified conditions, these solutions may converge to algorithms that are inefficient or erratic when applied to biological neuronal populations. It is, therefore, important to assess the performance of such algorithms with simple biological neuronal systems before moving to clinical trials. In this work, we tested, for the first time to the best of our knowledge, the efficacy of DFC in an *in vitro* setting. The many advantages of this method include the fact that it requires no prior knowledge about the system, imposes minimal perturbations and the actuation signal vanishes as the asynchronous state is approached.

Analogously to what was achieved *in silico*, we wanted to use DFC to disrupt the periodic bursting activity of biological neuronal networks. For that, we used randomly connected hippocampal networks that exhibited quasi-periodic synchronous bursting, making them suitable for the task.

### 3.1 Faster oscillations induced by non-adaptive DFC were not reported in *in silico* studies

Our implementation of non-adaptive DFC promoted a new bursting frequency instead of ablating it. This is in line with the claim that linear DFC could lead to synchronization if the delay deviated substantially from the half-period, T/2 (Popovych, Hauptmann, and Tass 2006). This was shown *in silico* by testing different delays with the same network. Here, however, the deviation from T/2 was caused by the fact that DFC stimulation induced a faster periodicity, while the delay was kept constant. The neuronal dynamics would align with the non-adaptive controller to the point that almost all activity was confined to stimulus responses (Figure S3). This was not predicted by computational studies, possibly because the simple *in silico* networks used in the aforementioned studies are not as dynamic as biological ones and, therefore, cannot sustain different synchronization periodicities. It is probably for this reason that this erratic behaviour of DFC was never observed *in silico*.

An important consideration is that the synchronization dynamics of the *in vitro* model are different from those explored in computational studies. Here, we were dealing with slow oscillations, in the order of seconds, whereas *in silico* studies implemented models whose oscillations were in the order of dozens of milliseconds. It is possible that these slow *in vitro* oscillations are more malleable by nature, allowing the network to sustain different periodicities. It remains to be explored whether DFC would produce the same erratic behaviour in biological networks with consistent fast oscillations. Having said that, there is nothing about the DFC method itself suggesting that it should only work for a certain range of periodicities. DFC requires no prior knowledge about the system and has already been implemented in many different (non-neurological) substrates, such as electrical oscillators, mechanical pendulums, electrochemical systems and others ^28^. In theory, DFC should have been a suitable candidate for a system that exhibits unstable periodic orbits, such as the quasi-periodic bursts of our *in vitro* networks.

### 3.2 Controllable networks may be at the vicinity of a state transition

We showed that only a subset of networks was differently modulated by each stimulation protocol. We refer to these as *controllable* networks. However, controllability is contingent on the experimental protocol. In particular, the fact that we only used one stimulating electrode may restrict the pool of controllable networks. We did so to follow the principle that DFC should control the system using minimal intervention (M. G. Rosenblum and Pikovsky 2004). Also, the choice of stimulating electrode and pulse amplitude plays an important role in the ability to drive the neuronal dynamics. In order to normalize this effect across cultures, we selected an electrode that elicited a network response using minimal pulse amplitude (see Methods for details). This way, we can consider each individual electrical pulse to be functionally similar for all networks. Differences in the susceptibility to modulation were, therefore, likely due to properties of the networks themselves. In order to explore this hypothesis, we plotted the spontaneous firing rate as a function of synchrony for all experiments. This showed that the baseline activity levels of controllable networks was somewhat unique, as the trials performed were clustered at the centre of the map, associated with intermediate levels of firing rate and synchrony.

The different arrangement of controllable and uncontrollable networks in the feature space may reflect the topology of the dynamical landscape of *in vitro* cultures. In fact, there is a vast literature exploring how some systems display particularly interesting properties when tuned to an intermediate level of a relevant order parameter. This sensitive point usually marks a phase transition between two distinct dynamical states, one typically characterized by too much order and the other by too much disorder. At the edge between states, these systems display optimal properties in terms of computation, information transmission and storage (Langton 1990). Some argue that neuronal systems can be included in such a class of systems and may even self-organize to meet the intermediate state where new interesting properties emerge (Shew et al. 2011; Beggs and Plenz 2003; Hesse and Gross 2014; Tetzlaff et al. 2010; Pasquale et al. 2008).

We postulate that the controllable networks may lie in a state transition associated with an intermediate level of some order parameter(s). However, we cannot conclude this simply by looking at the synchrony and firing rate as these are mere read-outs of the network activity, not parameters of the neuronal system. Using computer simulations, we verified that these read-outs could correlate with relevant network parameters, such as the excitatory/inhibitory balance and the overall synaptic strength of the neuronal connections. The parameter space established a transition from a state of too much disorder (random neuronal firing, too sparse to form oscillations) to a state of too much order (strong and highly synchronized activity). Most importantly, the networks that lay at the state transition were the ones exhibiting intermediate synchrony and firing rate and the ones where the controller actually worked; i.e., they were the controllable networks of this *in silico* analogy. This suggests that the controllable networks *in vitro* could have intermediate morphological features in terms of excitatory/inhibitory balance or strength of neuronal connections, for example.

We acknowledge that there are significant differences between the *in silico* and *in vitro* models, particularly regarding the time scales of the two systems. However, this scaling does not compromise the overall conclusion: the controllability region is associated with a state transition parametrized by specific network parameters, which generate intermediate levels synchrony and firing rate.

### 3.3 Asynchronous state is rarely reached: limitation of the algorithm or the *in vitro* model?

Despite the fact that aDFC decreased the synchrony of controllable networks, the neuronal activity was still dominated by synchronous events. These results were far from those obtained in *in* silico studies, where the synchronous events vanish almost completely. Among the multiple causes for that, we point to two possible sources: limitations of the algorithm implementation and limitations of the *in vitro* model itself.

Regarding the DFC algorithm, there was a practical constraint that limited its implementation in ideal conditions. Any controller should be capable of exerting positive and negative stimulation. However, with our setup, we only have access to excitatory electrical stimulation, much like existing implantable stimulators. For this reason, when the actuation signal (here modulated as a stimulation frequency) is negative, we send no input to the network. However, to implement DFC optimally, we should send inhibitory stimulation that follows the negative stimulation frequency. This could be achieved using optogenetics, which allows the concomitant use of both excitatory and inhibitory stimulation.

Nonetheless, in theory, using only the positive phase of the actuation signal should still prevent the formation of synchronous events, as it did in our *in silico* tests. So, it is possible that the limiting factor is the neuronal model itself. The unconstrained growth of our hippocampal cultures may have led to highly interconnected networks whose neurons simply cannot fire asynchronously. This is further supported by the fact that the only network where aDFC promoted the asynchronous state was one that could already sustain it autonomously. Again, these considerations are conditioned to the method used, whose main goal was to achieve desynchronization with minimal intervention, precisely timed at sensitive phases of the synchronization cycle. Other methods could possibly achieve network asynchrony by following more invasive approaches. Actually, it was already shown that using continuous high frequency stimulation (10-50 Hz, whereas our method uses around 1 Hz) distributed across multiple electrodes managed to prevent network synchronization, although at the cost of highly increasing the overall network firing rate (Wagenaar 2005).

There is still margin to improve the DFC algorithm by making it self-adaptive in terms of the actuation gain, pulse amplitude and selection of stimulation electrode(s), all of which were tuned manually before starting the experiments. It remains to be explored whether such an adaptive controller would consistently achieve full desynchronization in this type of network. However, it is a challenging problem to develop a controller with so many degrees of freedom, especially considering that it must operate in such a chaotic system. We also suggest other possible DFC implementations, already tested *in silico*, such as using a multi-electrode approach (Popovych and Tass 2018), modulating the actuation signal as pulse amplitude but keeping the stimulation frequency constant (Popovych et al. 2017) or using a continuous actuation signal instead of a pulsatile one (Vlachos et al. 2016), which would be particularly interesting with optogenetics.

To conclude, with the technological advances of implantable neurostimulators, there is an increasing need for algorithms capable of controlling pathological neuronal dynamics. Based on computational studies, DFC has been proposed as a promising candidate for disorders such as Parkinson’s disease and epilepsy. Our study marks an important milestone in the field by testing the efficacy of DFC in a biological context using microelectrode arrays to monitor and stimulate neuronal cultures. We showed that standard DFC is not only ineffective in ablating the oscillation, but actually worsens the oscillatory behaviour. We presented aDFC, which could disrupt the neuronal oscillations and reduce network synchrony. This result showed that it is vital for a DFC algorithm to follow the periodicity of the neuronal oscillations and adapt accordingly. We believe that the considerations extracted from this *in vitro* study will support the translation to future *in vivo* experiments.

## 5. Methods

### 5.1 Cell Cultures

The experiments followed both the European legislation regarding the use of animals for scientific purposes and the protocols approved by the ethical committee of i3S. The Animal Facility of i3S follows the FELASA guidelines and recommendations concerning laboratory animal welfare, complies with the European Guidelines (Directive 2010/63/EU) transposed to Portuguese legislation by Decreto-Lei no 113/2013 and is licensed by the Portuguese official veterinary department (DGAV, Ref 004461). Embryonic (E18) rat hippocampal neurons were cultured on 6-well or 9-well chamber MEA (256-6well MEA200/30iR-ITO-rcr and 256-9wellMEA300/30iR-ITO, respectively) (Multichannel System MCS, Germany) with a density of 5 × 10^4^ cells/well and 3 × 10^4^ cells/well, respectively. Each well of the 6-well MEA has 42 TiN electrodes (array of 7 × 6), and the 9-well contain arrays of 26 TiN electrodes (6 × 5 without corners). The experiments were performed between 13 and 55 days *in vitro*.

### 5.2 Electrophysiological Recordings

The recordings were performed with the MEA2100-256 System (Multichannel System, MCS, Germany). The cell cultures were kept at 37 °C and 5% CO2 using a stage-top incubator (ibidi GmbH, Germany) adapted to the headstage of the MEA2100-256 system. The electrophysiology signals were sampled at 10 kHz and filtered with a HP at 200 Hz (second order high pass butterworth filter). The neuronal spikes were detected and recorded using the Experimenter software from MCS. Peaks that were 6 STD (sometimes up to 8 STD, depending on the background noise) below or above baseline were considered neuronal spikes. There was a deadtime of 3 ms after each detection to prevent biphasic spikes from being identified twice. This spike data was used for posterior offline analysis.

### 5.3 Closed-loop Computer Application

The developed C# software was built using the McsUsbNet dynamic link library from MCS (https://github.com/multichannelsystems/McsUsbNet). The software ran in parallel with the Experimenter software in a quad-core Intel i7-6700 with 32.0 GB RAM workstation. The raw neuronal stream of data was received in batches at 10 kHz. Each batch had 1 sample point per electrode and was transferred directly from MEA2100-256 system via USB 3.0 high speed cable.

The signal processing was analogous to that running on Experimenter: the raw data was high-pass filtered at 200 Hz and the spike detection was performed using the same algorithm (section 5.2). The C# application had access to the MEA2100-256 System stimulators via USB 3.0, allowing us to trigger stimulation by software. The app also commanded the beginning of the recordings in Experimenter. The overall closed-loop latency associated with the activation of the stimulus was below 40 ms.

### 5.4 Delayed Feedback Control Algorithm

The C# application ran the DFC algorithm online. The application only monitored electrodes that had an average firing rate above of 0.1 Hz, measured before starting the experiments. The instantaneous firing rate (*FR*) of an active electrode was calculated in a square moving window of 0.1 s (sometimes tuned to 0.2 or 0.5 before starting the experiments depending on the dynamics of the network). The *FR* signal corresponds to the number of spikes per active electrode per unit of time. The *FR* of the monitored network was filtered using a damped oscillator ^17,18^,

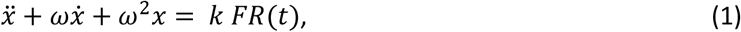

where *ω* denotes the natural frequency of the oscillator and *k* is a scaling coefficient. Here, we considered *k* = *ω*, which prevents the oscillator amplitude from scaling with the different ranges of periodicities exhibited by different cultures. We used 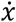 as the output of equation 1. The stimulation frequency (*SF*) is obtained by subtracting the current oscillation to the half-period shifted oscillation,

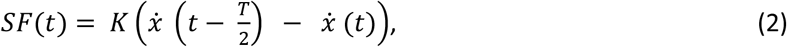

where *k* represents the stimulation gain and *T* is the oscillation period. A stimulus is triggered when

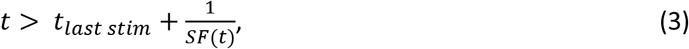

where *t* is the current time and *t*_*last stim*_ is the time stamp of the last stimulus. Equation 3 was bounded so that the stimulus was only sent when 1 < *SF*(*t*) < 20. The upper bound is required due to the latency in the stimulus activation (below 40 ms) and the lower bound guarantees that low background values of *SF* do not trigger stimulation (and that there is no negative stimulation frequency). Here, *T* and *ω* are adapted based on the real-time detection of synchronous events. The synchronous events (network bursts, NB) were detected when the *FR* signal was above a certain threshold, typically 10 Hz, considering a minimum interval between NB of 0.1 s. The threshold and minimum NB interval were sometimes adjusted before starting the experiments to better suit the observed data.

### 5.5 Experimental Protocol

Before starting the experiments, we monitored the networks for up to 10 min to identify the active electrodes. The application calculates the average firing rate for all the electrodes and selects those firing above 0.1 Hz. During this preliminary period, we also tuned, the parameters for the detection of synchronous events, if necessary.

Each stimulus unit was a negative electrical pulse with 200 µs duration. The choice of stimulation electrode and pulse amplitude was based on the premise that the stimulus should promote a response in the neighbouring electrodes using minimal stimulation. We probed different electrodes manually using Experimenter software, starting with the most active ones. For each electrode, we tested different stimulus amplitudes from -400 mV to -1000 mV in steps of -100 mV. When a given electrode and stimulation amplitude evoked a consistent response in nearby neurons, we fixed the stimulus amplitude to 120% of that value to account for eventual adaptations of the network, which could render the stimulation ineffective.

The experiments started with a recording of 15 min of spontaneous activity. The choice of stimulation protocols – DFC, aDFC and Poisson – for the following trials was randomized (with the exception noted below). Each trial had three periods: 5 mins without stimulation (OFF), 5 mins with stimulation (ON), and 5 mins without stimulation again (OFF). For the DFC and aDFC protocols, the stimulation followed the *SF*(*t*) signal determined by each algorithm. For the Poisson protocol, the stimulation followed a Poisson distribution of stimuli with an average frequency equivalent to that used in the previous aDFC trial (in the first round of experiments, the Poisson protocol was always the last of the three).

### 5.6 Selection criteria

We only performed experiments on networks that displayed consistent oscillatory bursting activity with periodicities between 0.5 and 5 seconds, approximately. Some cultures presented the required dynamics but were rejected because they were (or became) irresponsive to stimulation. In order to have statistical robustness in the evaluation of the different stimulation protocols, we only analysed networks that were submitted to at least three trials with each protocol. A total of 14 networks fulfilled the criteria to proceed with the experiments and analysis. Here, we define a network as being a neuronal culture in a given day *in vitro*. We did this because the neuronal dynamics of the same culture of neurons may change drastically from day to day, so we considered it a different dynamical system. The 14 networks came from 11 independent cultures, from a total of five different animal preparations (Figure S2).

### 5.7 Signal processing

#### Features of neuronal activity

The offline analysis of neuronal data was based on the spikes recorded with Experimenter. We used four features to characterize the activity of the different networks: average firing rate, synchrony, main oscillation frequency and intensity. The average firing rate corresponds to the average number of spikes per electrode per unit of time. The synchrony measure is based on the *χ* metric, which compares the variance of the network activity with the variance of each neuron’s activity ^29^,

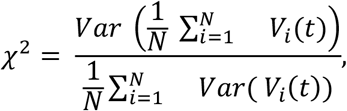

where *N* is typically the number neurons, but here corresponds to the number of electrodes, and *V*_*i*_(t) is typically the voltage of neuron *i*, but here corresponds to the spike train of electrode *i* convolved with a square wave of 50 ms and binarized. This signal defines the periods on which a given electrode is considered active. We are thus measuring the degree of coactivation of the multiple electrodes, regardless of the number spikes exhibited by different electrodes at the moment of synchronization. We did not want this factor to leak to the synchrony measure because different electrodes may be synchronized but with different number of spikes per activation (which could be due to different number of neurons surrounding each electrode, for example).

The main oscillation frequency and intensity was obtained by identifying the peak (x and y coordinates, respectively) of the discrete Fourier transform of the instantaneous firing rate of the entire network. The instantaneous firing rate of the network was obtained by convolving the spike train of each electrode with a Gaussian kernel of 10 ms and averaging across electrodes.

#### Unsupervised clustering of neuronal dynamics

The different states of neuronal activity of a given recording (section 2.5) were identified by calculating the firing rate and synchrony in a moving window. The width of the moving window corresponded to the average duration of five periodic events in the analysed recording. The moving step was half the window width. The clusters of neuronal dynamics were automatically identified with Gaussian Mixture Models (GMMs) applied to the bidimensional matrix of firing rate and synchrony. We tested GMMs with 1 to 4 groups and each condition was repeated 10 times with different initial values. The ideal clustering was chosen based on the Bayesian Informative Criterion. We calculated the centroid of each cluster and considered the asynchronous states as being those with mean synchrony below 0.5 and mean firing rate below the average for that network.

### 5.8 Statistical Analysis

#### Controllable and uncontrollable networks

The classification of networks as controllable or uncontrollable (Figure 3) was based on the assumption that controllable networks could be driven to specific subspaces of neuronal activity with a given stimulation protocol whereas the uncontrollable networks could not. We started by calculating the fold change of the four metrics (synchrony, firing rate, oscillation frequency and intensity) for all trials performed with each network. To evaluate whether the spaces reached during stimulation were statistically different, we calculated the Multivariate ANOVA (MANOVA) considering the fold change of the four metrics, simultaneously. Here, the null hypothesis was that the modulation results were identical for all stimulation protocols.

#### Neuromodulation results for controllable networks

For a given network, we calculated the average fold change of each metric across all the trials performed with each stimulation protocol. Then, comparisons between control and test groups (aDFC, DFC and Poisson) were performed using two-tailed paired t-tests.

#### Time spent in asynchronous state

To compare the efficacy of the different stimulation protocols in promoting the transition to a stable asynchronous state in a multi-stable network, we calculated the percentage of time spent in the asynchronous state during the OFF and ON periods for all the trials. We then compared if there was a significant change from OFF pre-stimulation to ON and from OFF pre-stimulation to OFF post-stimulation using two-tailed paired t-tests.

### 5.9 In silico model

Our *in silico* networks were composed of 1000 randomly connected Izhikevich neurons with varying percentages of regular spiking excitatory neurons (a = 0.02; b = 0.2; c = -65; d = 8) and fast spiking inhibitory neurons (a = 0.1; b = 0.2; c = -65; d = 2) ^27^. The neurons were connected in an all-to-all architecture of varying synaptic weights according to the connectivity matrix S. The overall synaptic weights of the different simulations were modulated by scaling S. Each condition was simulated five times. The model was based on the script provided in <https://www.izhikevich.org/publications/net.m.> The simulations had an initial period of 500 ms for the network activity to stabilize. After the stabilization period, we had 2000 ms of spontaneous activity and 2000 ms under the aDFC stimulation protocol. The aDFC algorithm was identical to that applied *in vitro*, considering only the positive component of the actuation signal. A pool of 100 neurons was randomly selected for stimulation. Each electrical pulse corresponded to an instantaneous input of 20 mA to the stimulated neurons. The firing rate and synchrony measures were computed using the same algorithms as those used *in vitro* (section 5.7) and averaged across the five repetitions of each simulation condition.

## Acknowledgements

This work was partially financed by FEDER—Fundo Europeu de Desenvolvimento Regional funds through the COMPETE 2020—Operacional Programme for Competitiveness and Internationalisation (POCI), Portugal 2020, and by Portuguese funds through FCT—Fundação para a Ciência e a Tecnologia/Ministério da Ciência, Tecnologia e Ensino Superior, in the framework of project PTDC/EMD-EMD/31540/2017 (POCI01-0145-FEDER-031540). Domingos Leite de Castro was funded by FCT – Fundação para a Ciência e a Tecnologia, grant contract SFRH/BD/143956/2019.

## Author contributions

D.C. contributed to methodology, experiments, software, formal analysis, visualization, and writing of original draft. M.A. contributed to experiments and formal analysis. A.P.A. and D.G. contributed to formal analysis. P.A. contributed to conceptualization, methodology, funding, and work supervision. All authors contributed to the review and editing of the final manuscript.

## Data Availability

The data that support the findings of this study are available from the corresponding author upon reasonable request.

## Supplementary Information

**Supplementary Figure S1.**
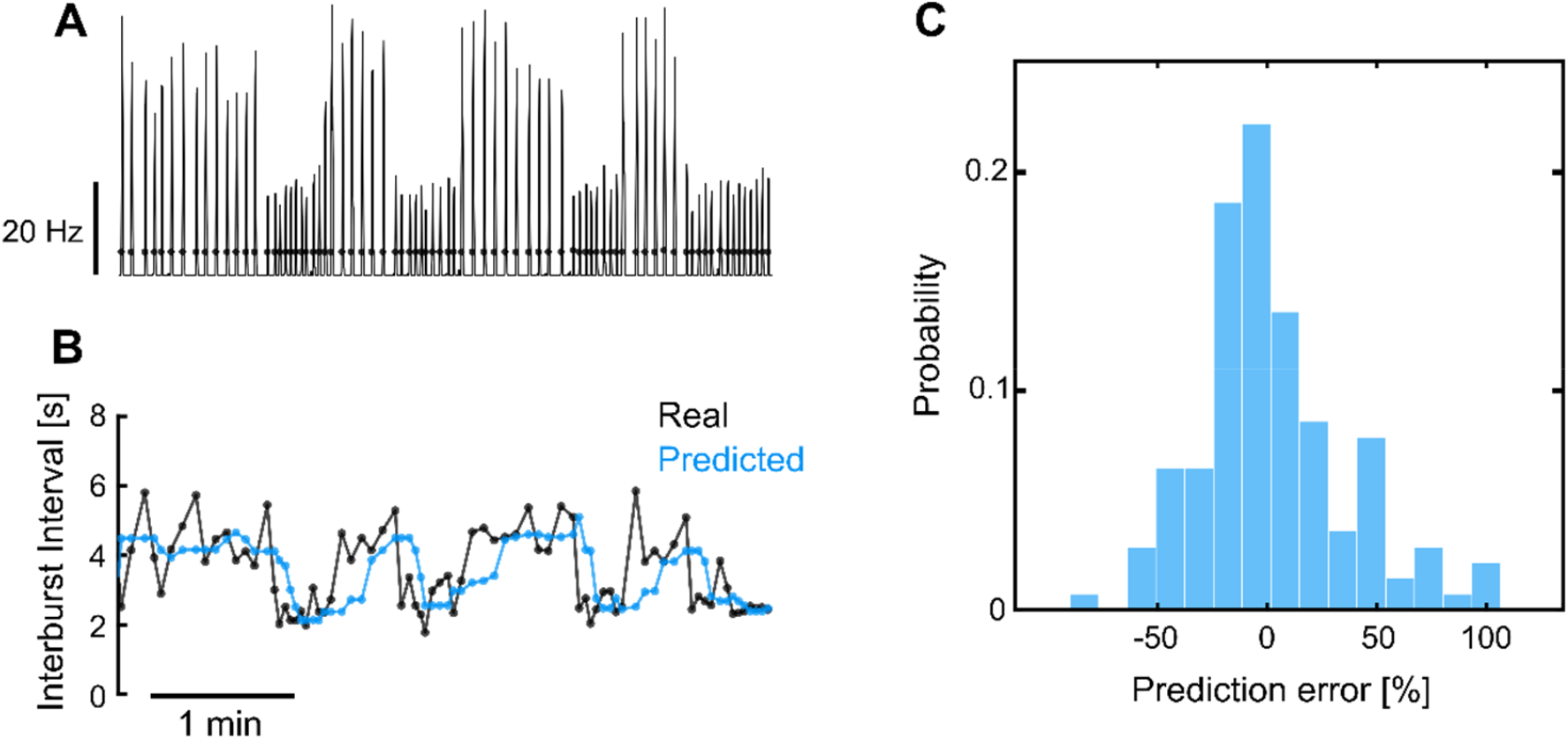
Representative example of aDFC adaptability in a network with fast changing dynamics. A) Instantaneous network firing rate during spontaneous activity. The black dots represent the real-time detection of network bursts performed by the adaptive controller. B) Interburst interval predicted by the aDFC controller and the associated real interbursts. Every time the controller detects a new network burst (black dots in A) it estimates when the next burst would spontaneously occur by calculating the median of the previous five interburst intervals. This prediction is used to properly tune the controller’s half-cycle delay and the natural frequency of the oscillators used to create the actuation signal. In this example the stimulation is turned OFF, so the controller is simply tracking the neuronal activity without interfering with it. C) Histogram of the prediction error for the representative example shown in A and B. The largest errors result from the prediction lag associated with the fast transitions between bursting regimes. The fact that the prediction error is typically small, even under this fast changing network dynamics, means that the half-cycle delay and the oscillator’s natural frequency of aDFC are properly self-tuned in real-time.

**Supplementary Figure S2.**
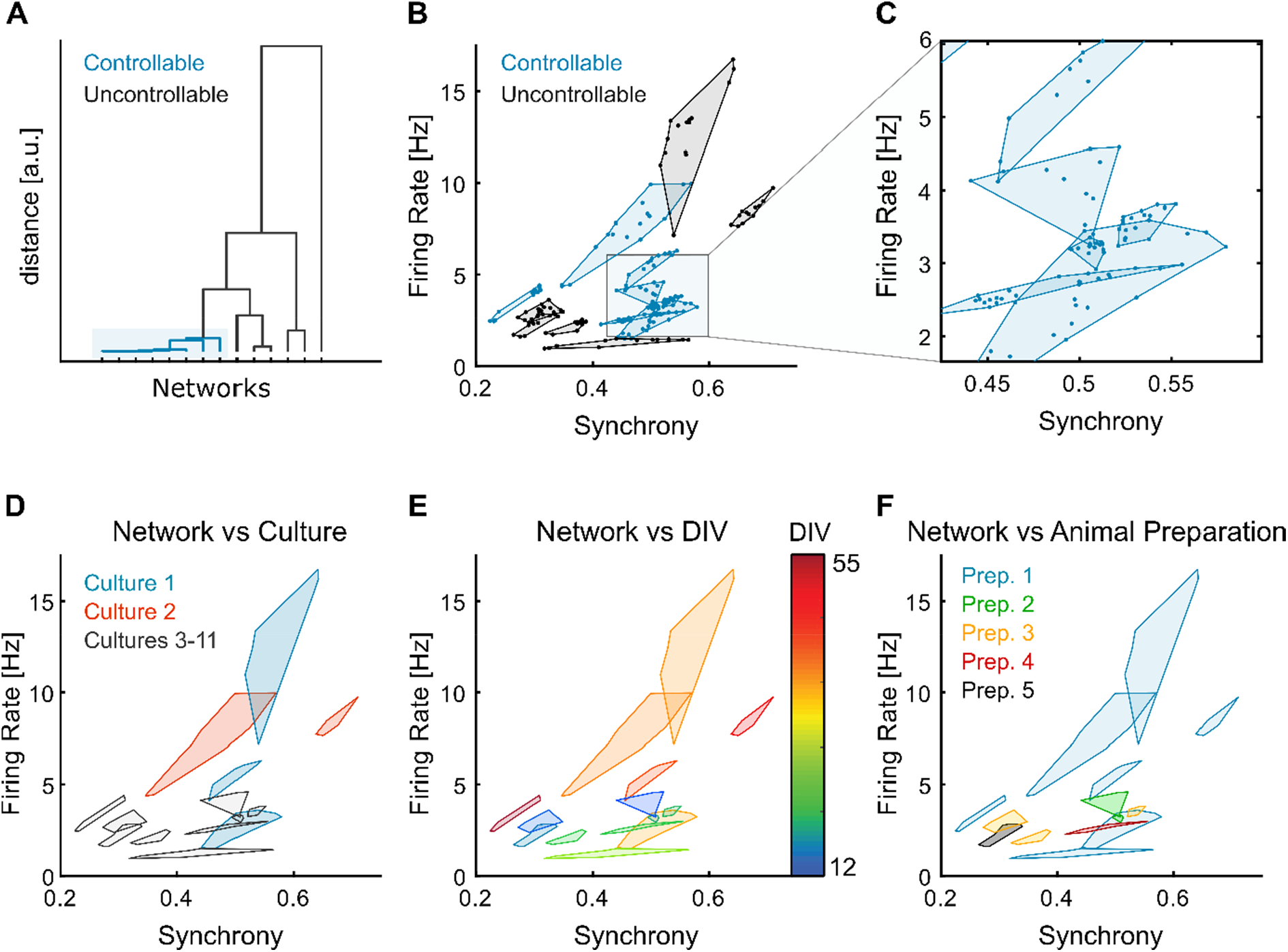
Controllable and uncontrollable networks across cultures, DIVs and animal preparations. A) Dendrogram of the networks according to the controllability measure (MANOVA p-values of the modulation results obtained with the different stimulation protocols). This representation shows that the controllable networks (p-values < 0.05) are more similar to each other than to the uncontrollable ones. B) Spontaneous firing rate and synchrony for the all the trials of the different networks, separated into controllable and uncontrollable networks. C) Detail of B, evidencing how some cultures have very well confined spontaneous dynamics. D) Division of the networks according to the associated neuronal culture. When probed in different days, the same neuronal culture can be considered a different dynamical system because the spontaneous dynamics are clearly different. Therefore, we consider these to be different networks. For example, culture 1 (blue) and culture 2 (orange) led to three and two different networks, respectively, some being controllable and some uncontrollable for the same culture. Cultures 3 to 11 (grey) only have one network each and are represented with the same color for clarity. E) Evolution of the networks with the DIV. This representation suggests that there is a trend towards higher levels of firing rate and synchrony with the DIVs. Also, the DIV by itself does not predict which networks are controllable and uncontrollable as these groups span multiple overlapping DIVs. F) The networks used came from a total of five different animal preparations. The distinction of controllable and uncontrollable networks cannot be attributed to any particular preparation, as the same preparation can lead to both types of networks.

**Supplementary Figure S3.**
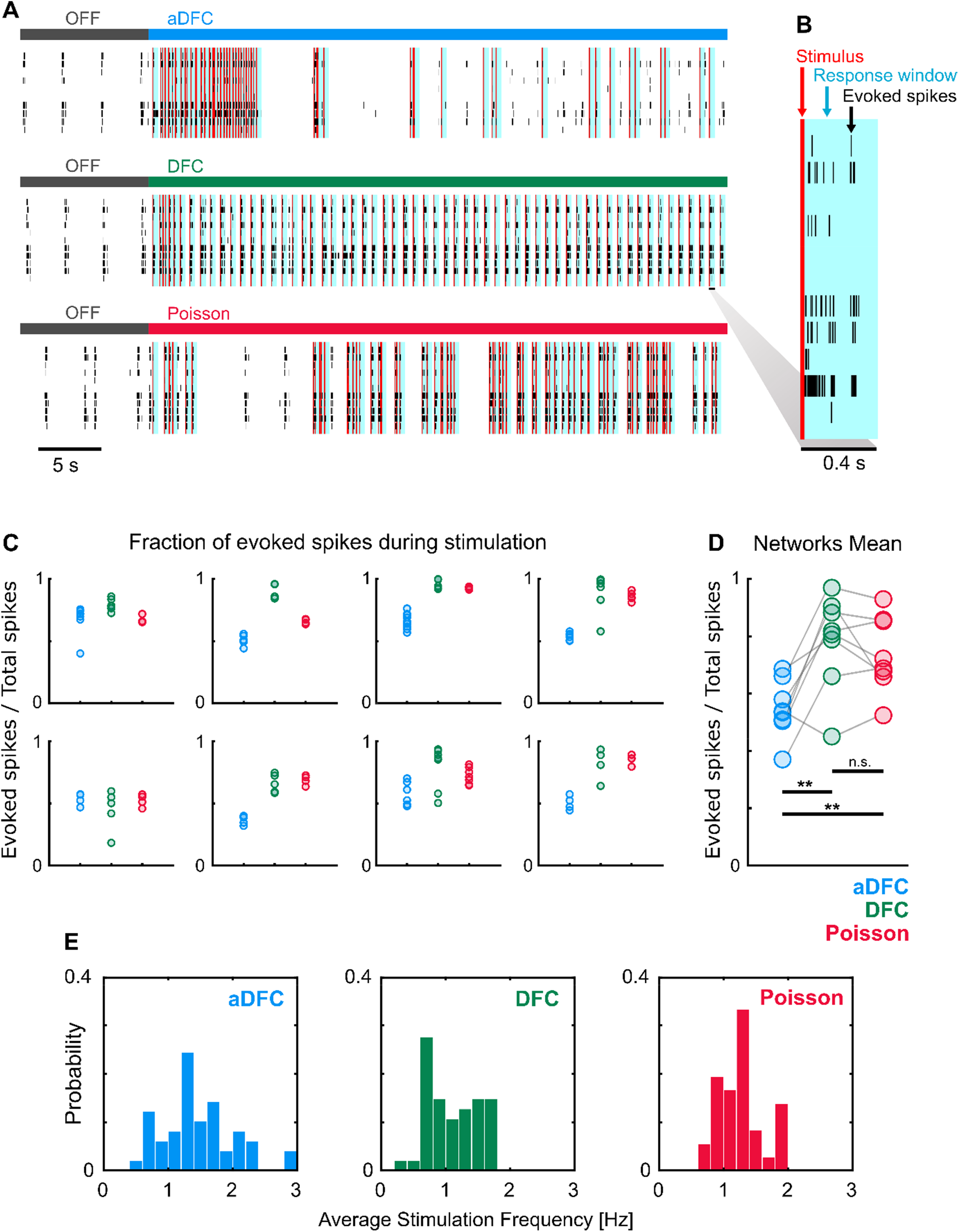
aDFC allows the network to maintain some autonomous firing whereas DFC and Poisson tend to confine most network activity to stimulus responses. A) Stimuli (red) and respective response windows of 400 ms (blue) for representative examples of aDFC (top), DFC (middle) and Poisson (bottom) applied to the same network. B) Detail of the spikes (black) evoked after stimulation (red) inside the response window (blue). C) Fraction of evoked spikes during stimulation for all the trials performed with the three protocols for each controllable network (one plot per network). When the fraction of evoked spikes is close to 1, the network has almost no spontaneous activity and is simply replicating the actuation signal, which is not ideal. D) Comparison of the mean fraction of evoked spikes for the controllable networks. Each triplet of point corresponds to one network. aDFC has a significantly lower percentage of evoked activity, meaning that it allows the network to maintain some intrinsic activity. The statistical tests consisted in t-tests (n = 8; **, p < 0.01; details of statistical tests in Table S3). E) Average stimulation frequency used by each method for all trials performed with all networks. The overall stimulation frequency is within the same range for all methods. This shows that the results found in D are not explained by differences in the amount of stimulation but are rather due to the specific timing of the stimuli determined by each algorithm.

**Supplementary Table S1.** Details of the statistical tests used in Figure 4 to compare the modulation results of the different stimulation protocols in controllable networks (two tailed ratio paired t-tests)

|  | Oscillation Intensity |  |  | Oscillation Frequency |  |  |
| --- | --- | --- | --- | --- | --- | --- |
|  | aDFC | DFC | Poisson | aDFC | DFC | Poisson |
| P value | 0.0011 | 0.0248 | 0.0123 | 0.0608 | 0.0027 | 0.0795 |
| t | 5.279 | 2.847 | 3.346 | 2.232 | 4.523 | 2.050 |
| deg. f. | 7 | 7 | 7 | 7 | 7 | 7 |
| n° pairs | 8 | 8 | 8 | 8 | 8 | 8 |

|  | Synchrony |  |  | Firing Rate |  |  |
| --- | --- | --- | --- | --- | --- | --- |
|  | aDFC | DFC | Poisson | aDFC | DFC | Poisson |
| P value | 0.0418 | 0.1057 | 0.1283 | 0.0021 | 0.0481 | 0.0434 |
| t | 2.487 | 1.857 | 1.724 | 4.731 | 2.391 | 2.461 |
| deg. f. | 7 | 7 | 7 | 7 | 7 | 7 |
| n° pairs | 8 | 8 | 8 | 8 | 8 | 8 |

**Supplementary Table S2.** Details of the statistical tests used in Figure 7 to compare percentage of time spent in asynchronous state for each stimulation protocol in a multistable network (two tailed paired t-tests)

|  | aDFC |  |  |
| --- | --- | --- | --- |
|  | OFF pre vs ON | ON vs OFF pos | OFF pre vs OFF pos |
| P value | <0.0001 | <0.0001 | 0.7281 |
| t | 11.3 | 9.995 | 0.3587 |
| df | 9 | 9 | 9 |
| n° pairs | 10 | 10 | 10 |

|  | DFC |  |  |
| --- | --- | --- | --- |
|  | OFF pre vs ON | ON vs OFF pos | OFF pre vs OFF pos |
| P value | 0.5088 | 0.1598 | 0.0808 |
| t | 0.7024 | 1.604 | 2.097 |
| df | 6 | 6 | 6 |
| n° pairs | 7 | 7 | 7 |

|  | Poisson |  |  |
| --- | --- | --- | --- |
|  | OFF pre vs ON | ON vs OFF pos | OFF pre vs OFF pos |
| P value | 0.0935 | 0.3909 | 0.5229 |
| t | 2.428 | 1 | 0.7211 |
| df | 3 | 3 | 3 |
| n° pairs | 4 | 4 | 4 |

**Supplementary Table S3.** Details of the statistical tests used in Figure S3 to compare fraction of spontaneous spikes during stimulation for the three stimulation protocols (two tailed paired t-tests)

|  | aDFC vs DFC | aDFC vs Poisson | DFC vs Poisson |
| --- | --- | --- | --- |
| P value | 0.0037 | 0.0075 | 0.2394 |
| t | 4.281 | 3.712 | 1.286 |
| deg. f. | 7 | 7 | 7 |
| nº pairs | 8 | 8 | 8 |

